# Comparing the functional structure of neural networks from representational similarity analysis with those from functional connectivity and univariate analyses

**DOI:** 10.1101/487199

**Authors:** Ineke Pillet, Hans Op de Beeck, Haemy Lee Masson

## Abstract

The invention of representational similarity analysis (RSA, following multi-voxel pattern analysis (MVPA)) has allowed cognitive neuroscientists to identify the representational structure of multiple brain regions, moving beyond functional localization. By comparing these structures, cognitive neuroscientists can characterize how brain areas form functional networks. Univariate analysis (UNIVAR) and functional connectivity analysis (FCA) are two other popular methods to identify the functional structure of brain networks. Despite their popularity, few studies have examined the relationship between the structure of the networks from RSA with those from UNIVAR and FCA. Thus, the aim of the current study is to examine the similarities between neural networks derived from RSA with those from UNIVAR and FCA to explore how these methods relate to each other. We analyzed the data of a previously published study with the three methods and compared the results by performing (partial) correlation and multiple regression analysis. Our findings reveal that neural networks resulting from RSA, UNIVAR, and FCA methods are highly similar to each other even after ruling out the effect of anatomical proximity between the network nodes. Nevertheless, the neural network from each method shows idiosyncratic structure that cannot be explained by any of the other methods. Thus, we conclude that the RSA, UNIVAR and FCA methods provide similar but not identical information on how brain regions are organized in functional networks.

## 1. Introduction

Multi-voxel pattern analysis (MVPA) has recently become one of the most frequently used techniques for analyzing fMRI data. It considers the spatial pattern of neural activation across multiple voxels and examines whether these patterns contain task-related information (Coutanche, 2013; Haxby, 2012; Haxby et al., 2001; Haynes, 2015; Haynes and Rees, 2006; Kriegeskorte, 2011; Mur et al., 2009). It is referred to as *multivariate* or *multi-voxel* because it analyzes a set of voxels together (the pattern of activation of this set) instead of modeling activity of a single voxel (as is done in univariate analysis) (Kriegeskorte, 2011; Mur et al., 2009; Norman et al., 2006; Yang et al., 2012). In addition, patterns of activation can be used to investigate the similarities between such patterns of different conditions, or between such patterns of different brain regions in a certain condition (Haxby, 2012, Mur et al., 2009). This approach is referred to as representational similarity analysis (RSA) (Kriegeskorte et al., 2008). In first-order RSA, a representational dissimilarity matrix (RDM) is set up to understand the dissimilarity between patterns of activation of different stimuli in a certain brain region (Kriegeskorte et al., 2008; Yang et al., 2012). In second-order RSA, RDMs are compared between brain regions (Kriegeskorte et al., 2008; Yang et al., 2012). This method has been referred to as *representational connectivity* as it allows to identify the representational relationship among brain regions (Kriegeskorte et al., 2008). Connectivity related to multivariate information has since then been given a more specific meaning to refer to analyses of the temporal dynamics of the information contained in multi-voxel patterns, also sometimes referred to as multivariate or informational connectivity (Anzellotti and Coutanche, 2018; Coutanche and Thompson-Schill, 2014). For this reason, we opted for the more general RSA term instead of using the term representational connectivity.

Univariate analysis (UNIVAR) and functional connectivity analysis (FCA) are two other frequently used techniques for analyzing fMRI data. UNIVAR assesses neural activation of an individual voxel or a mean activation across voxels of a brain region. For this reason, it is often used to localize brain regions engaged in processing a particular type of stimuli (e.g., face versus object) and thereby draw conclusions about the regions that are involved in cognitive processes important for the stimuli or task at hand (Coutanche, 2013; Haynes, 2015; Haynes and Rees, 2006; Logothetis, 2008; Mur et al., 2009). It is referred to as *univariate* because a general linear model (GLM) is applied voxel-wise to relate the experimental design to the neural activity of each voxel’s time-course in the brain (Raizada and Kriegeskorte, 2010). FCA (for a review, see Friston, 2011) characterizes the communication between brain regions during rest or a task (Friston, 1994), measuring the strength of the relation between BOLD time-series signals of brain regions (Geerligs et al., 2016; Yang et al., 2012). When FCA is applied to a resting-state fMRI dataset, it reveals the intrinsic network of the brain based on low-frequency BOLD fluctuations of brain regions (Biswal et al. 1995; Cordes et al., 2001; Fox and Raichle, 2007). This intrinsic network can also be extracted from a task-based fMRI dataset by removing the task-induced signal from the data (Fair et al., 2007). Therefore, this method is often referred to as *intrinsic functional connectivity*.

Notably, there are various conceptual similarities and differences between RSA, UNIVAR and FCA. UNIVAR and FCA methods are similar in that they average BOLD signal of the voxels in the brain region, unlike MVPA. The structure of networks from co-activation has also proven to be similar to those from resting-state connectivity (Crossley et al., 2013). Analogously, Anzellotti and Coutanche referred to this type of FCA as univariate FCA (Anzellotti and Coutanche, 2018). Second-order RSA and FCA are similar in that they are both based on a measure of the similarity between brain regions. When using correlations, correlating the averaged BOLD time-series signals between the ROIs in FCA is methodologically similar to correlating RDMs of those ROIs in second-order RSA (Xue et al., 2013). UNIVAR and RSA, or at least MVPA, have been frequently compared when describing functional properties of one region of the brain (e.g., see Coutanche, 2013; Davis et al., 2014; Gilron et al., 2017; Jimura and Poldrack, 2012). A significant finding from these studies was that changes (across different stimuli) in the activation patterns could be detected even when conditions were not different in the average univariate activation in a region (Mur et al., 2009). For example, different speech sounds showed different activation patterns in the right auditory cortex, but the average-activation of this region across those speech sounds did not differ (Raizada et al., 2010). These studies have provided valuable insights into the conceptual and empirical relationships between UNIVAR and MVPA. Similarly, studies have used both RSA and FCA, some drawing the same conclusion from the results of RSA and FCA (e.g., Zeharia et al., 2015), or not (Boets et al., 2013; Bulthé et al., 2018).

A study that directly and simultaneously compares network structures resulting from RSA with those from UNIVAR and FCA is missing from today’s literature. Thus, the current study explores how the networks from RSA complement the networks from UNIVAR and FCA when investigating the functional architecture of the brain. Although direct and simultaneous comparisons between brain network structures based on UNIVAR, RSA and FCA have not been performed (to our knowledge), we expect at least some convergence. For example, we hypothesize that brain regions with similar representational similarity structure would tend to be functionally connected, without excluding the possibility of uniqueness in the networks resulting from the two methods. Specifically, given the evidence of the topographic arrangement of the basic sensory cortical areas, such as the visual and sensorimotor cortex (see Kaas, 1997, for a detailed review), we predict that the way in which brain networks composed of visual or sensorimotor areas are constructed would be highly similar in all three methods. In sum, the goal of this study is to explore how the network structure derived from RSA compares to those from UNIVAR and FCA. To answer this question, we applied second-order RSA to a previously reported fMRI study (Lee Masson et al., 2018) and compared the resulting networks with the results obtained from UNIVAR and intrinsic FCA. In particular, we conducted (partial) correlation and multiple regression analysis (comparing them to signal-to-noise ratio measurements), controlling for the confounding influence of anatomical proximity between brain regions of interest on RSA, UNIVAR and intrinsic FCA results. In addition, we explored our results visually by implementing multi-dimensional scaling (MDS) and Procrustes transformation methods.

## 2. Methods

### 2.1 Datasets

We reanalyzed data from our previous fMRI study (Lee Masson et al., 2018). All participants provided written informed consent before the experiment in accordance with the Declaration of Helsinki. The study was carried out in accordance with the recommendations of and 12approved by the Medical Ethical Committee of KU Leuven (S53768 and S59577). In this study, 21 healthy participants observed greyscale videos (see Fig. 1) of social touch interaction, varying in valence and arousal (Lee Masson and Op de Beeck, 2018). The experiment included 39 social touch videos, and in addition 36 nonsocial control videos. Participants carried out an orthogonal attention task: they pressed a button with their left or right thumb whenever the touch interaction initiator wore a grey or black shirt, depending on the instruction of that specific run (left for grey, right for black). The stimuli were displayed for 3 s, followed by an inter-stimulus interval of 3 s during which a fixation cross was presented and during which the participants could press a button as a response related to the task. Each run was divided into three blocks of 25 videos. At the start of each block, a baseline (display of a fixation cross) of 6 s was included. The total duration of each run was 7.80 min. The participants completed six runs. In the following UNIVAR and RSA analyses, we restrict the analyses to the data from the 39 social touch videos.

**Fig. 1.**
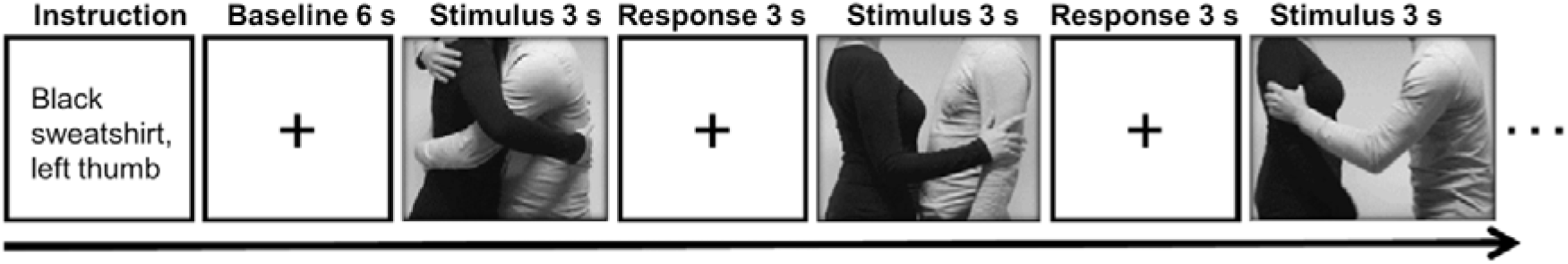
The experimental procedure. Participants received an instruction on when they should press a certain button (e.g. press the button with your left thumb when touch interaction initiator wears black sweatshirt). After a baseline of 6 s, the stimuli were presented for 3 s always followed by an inter-stimulus interval of 3 s, during which a fixation cross was presented and participants could press a button. In this example, still frames of three social touch videos are shown (left: hug, middle: stroke, right: shake). All videos can be found here: https://osf.io/nq5mf/

Importantly, when creating the videos, we controlled for the visual elements, such as clothes style and color of the actors so that these do not induce a visually biased neural response (Lee Masson and Op de Beeck, 2018). For example, having actors wear blue in the pleasant touch scenes and having actors wear red in unpleasant touch scenes can induce visual bias related to the clothing color when contrasting the brain response between the pleasant and unpleasant touch conditions.

In addition, the scan sessions included runs in which participants received (instead of observing) pleasant (brush strokes) and unpleasant touch (rubber band snaps) in a block design (see Lee Masson et al., 2018, for more details). These data were used for the intrinsic FCA.

### 2.2 Regions of interest (ROIs)

For our previous study (Lee Masson et al., 2018), we selected 16 a priori defined ROIs, belonging to four different networks in the brain that proved to be important in processing observed social touch interactions: the somatosensory-motor network (the parietal operculum (PO), Brodmann area (BA) 3, BA1, BA2, BA4 (Rolls et al., 2003)), the social-cognitive network (the middle temporal gyrus (MTG), the precuneus, the superior temporal gyrus (STG), the temporoparietal junction (TPJ) (Jacoby et al., 2016)), the pain network (the middle cingulate cortex (MCC), the insula (Gordon et al., 2013; Lamm and Majdandžić, 2015; Morrison et al., 2011), and the visual network given that visual stimuli were used (BA17, BA18, BA19, BA37, V5) (see Fig. 2). To define these ROIs anatomically, first, we made masks with various templates from PickAtlas software (Maldjian et al., 2003), SPM Anatomy toolbox (Eickhoff et al., 2005) and connectivity-based parcellation atlas (Mars et al., 2012). Second, we extracted all the voxels in the mask per ROI and combined left and right hemispheres. Afterwards, we examined if there were overlapping voxels among ROIs (e.g., V5 is located in BA19 and BA37) and removed overlapping voxels from each other in order to ensure all ROIs are anatomically independent. For further information about these ROIs and how they were defined, see our previous study (Lee Masson et al., 2018). In contrast to FCA that often includes a more extensive set of ROIs, the RSA method requires ROIs to contain meaningful neural signals associated with the experimental conditions. For this reason, only the aforementioned 16 ROIs, whose spatial neural patterns passed the MVPA reliability test, were selected (Lee Masson et al., 2018). Briefly, in this reliability test, runs are split into two halves and the correlation between neural patterns for within- and between-conditions are compared per ROI. This process is repeated 100 times (to randomly split the runs into two halves) and these results are then averaged. ROIs are only included if the correlations for within-condition comparisons are significantly stronger than those for between-condition comparisons. Neural pattern similarity between different conditions most likely only reflect noise when neural pattern similarity between the same conditions is low (Ritchie et al., 2017; for more details on this test and the results see Lee Masson et al., 2018).

**Fig. 2.**
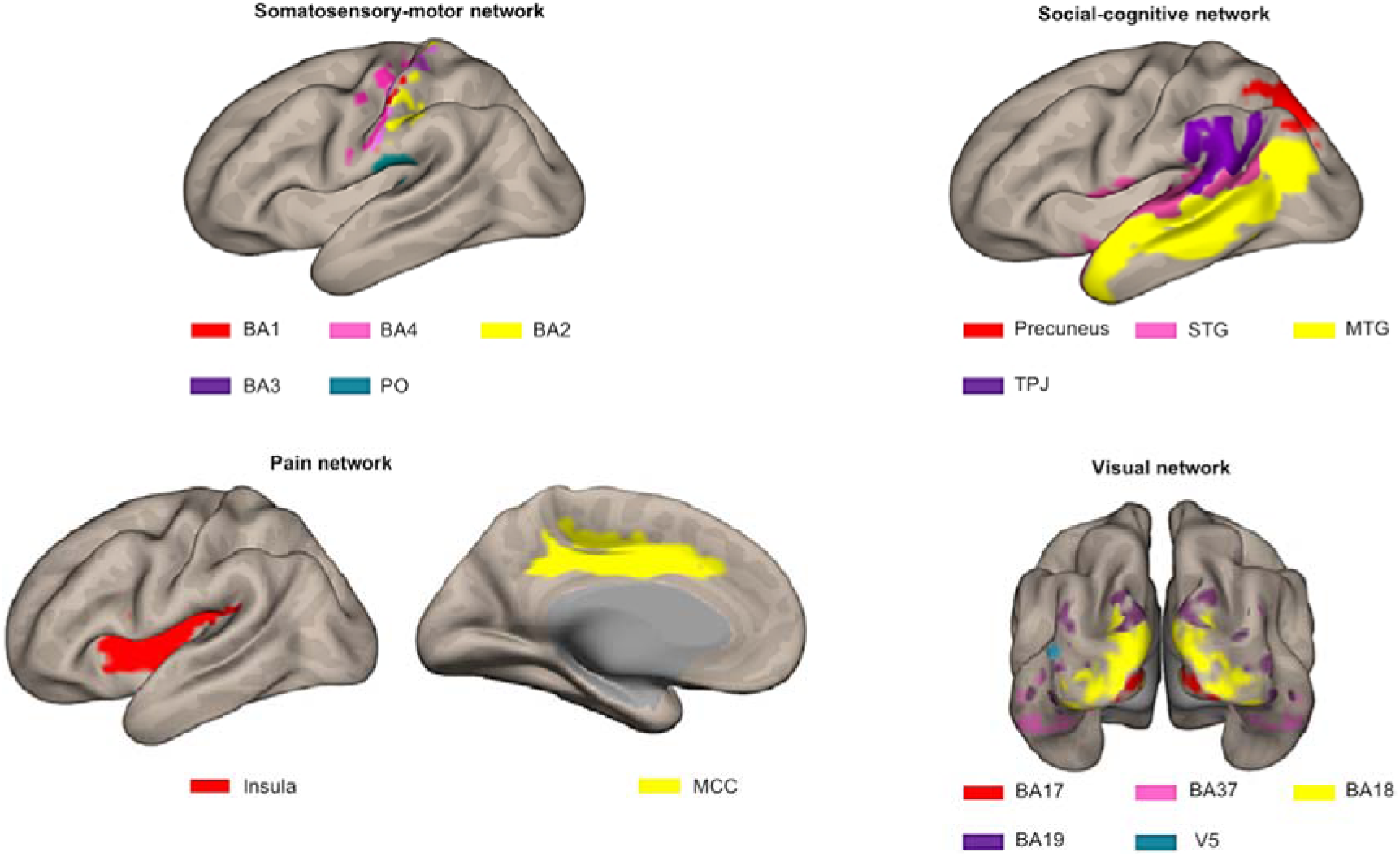
Illustration of the different ROIs in the context of the network they were assigned to a priori. Top left: somatosensory-motor network ROIs including BA1 (red), BA4 (pink), BA2 (yellow), BA3 (purple) and PO (blue). Top right: social-cognitive network ROIs including precuneus (red), STG (pink), MTG (yellow) and TPJ (purple). Bottom left: pain network ROIs including insula (red) and MCC (yellow). Bottom right: visual network ROIs including BA17 (red), BA37 (pink), BA18 (yellow), BA19 (purple) and V5 (blue). This figure was made using CONN toolbox 17 (Whitfield-Gabrieli and Nieto-Castanon, 2012)

### 2.3 Univariate analysis

In our previous study, we processed functional data by using a standard preprocessing pipeline and by applying a general linear model (GLM) to each subject’s data (Lee Masson et al., 2018). On top of the regressors of interest (matched to the onset time of each regressor (duration = 0) of the event-related design of the fMRI observing touch experiment), six head motion parameters were included in the models as nuisance covariates (Lee Masson et al., 2018). These GLMs were defined with data smoothed with 8 mm FWHM. For detailed information on how data was preprocessed and how the GLM was applied, see our previous study (Lee Masson et al., 2018). From these GLMs, we obtained the estimated beta-values per stimulus of the social condition (N = 39) for all voxels in each ROI. For each stimulus, we averaged the beta values of all runs, of all voxels within each ROI, and of all participants, yielding a one-dimensional array with 39 elements in each ROI, reflecting how strongly each of the 39 videos activated the ROI. These arrays were Pearson correlated for each possible pair combination of ROIs to investigate the similarity between ROIs’ average BOLD responses evoked during the observation of social touch and therefore to investigate clustering/networks of our ROIs with regard to their average activations. We refer to this clustering as the activation network.

### 2.4 Representational similarity analysis

This analysis was based upon a GLMs applied to the fMRI observing touch experiment that consisted of 75 predictors (one for each video). The preprocessing pipeline for this analysis differed slightly from the one used for univariate analysis: data was smoothed at 5 mm FWHM. As such, we optimized the preprocessing parameters to fit the requirements of each analysis (Hendriks et al., 2017). For each ROI, we created a 39 × 39 neural matrix by correlating (Pearson) the multi-voxel patterns between all possible combinations of pairs of stimuli of the social condition (*N* = 39) and then averaged this across subjects (first-order RSA, Kriegeskorte et al., 2008). After, we vectorized the upper diagonal elements of this group-averaged matrix while discarding the diagonal and lower diagonal elements, and correlated (Pearson) these vectors for all possible combinations of ROI pairs (second-order RSA, Kriegeskorte et al., 2008). These comparisons between areas allow us to investigate the representational similarity between ROIs and therefore to investigate clustering/networks of our ROIs with regard to the between-condition similarity in multi-voxel activation patterns (Kriegeskorte et al., 2008). We refer to this clustering as the representation network. More information on the details of how MVPA was applied to fMRI data can be found in our previous study (Lee Masson et al., 2018).

### 2.5 Functional connectivity analysis

Functional connectivity analysis, performed in the CONN toolbox 17 (Whitfield-Gabrieli and Nieto-Castanon, 2012), was applied to a different set of fMRI data (wherein participants received touch) obtained in the same scan sessions. We used two independent sets of fMRI data to avoid a spurious correlation between the two sets of brain networks resulting from UNIVAR and FCA. BOLD signal fluctuations may be partially induced by the presented stimuli, which may result in shared signals between networks derived from the UNIVAR and FCA methods. In the other direction, spontaneous fluctuations over time may affect the estimated univariate activation.

Preprocessing was conducted as described in our previous study (Lee Masson et al., 2018), but again optimized to fit the requirements of FCA: no smoothing was carried out to avoid a spillover effect (Alakörkkö et al., 2017). The outlier scans were detected based on the global signal spike and motion in the functional data by the Artifact Detection Toolbox (ART) software package (www.nitrc.org/projects/artifact_detect/). Consequently, standard denoising methods were applied to remove confounding effects. This step consists of 1) linearly regressing out 13 principal components of white matter and cerebrospinal fluid signals, six head motion parameters and their first-order derivatives, all scrubbing covariates from the artifact detection, and main task effects (rest condition, see below), 2) linear detrending, and 3) band-pass filtering (0.008-0.09 Hz) that removes slowly fluctuating noise, such as scanner drift, and the task-induced signal. To calculate intrinsic FC (functional connectivity), we did not encode task-related information in the experimental design. Instead, task effect (i.e., receiving touch) was removed from the fMRI time series by including regressors corresponding to each task condition during denoising step, and the rest condition was defined (Fair et al., 2007). Previous studies indicated that the intrinsic fluctuations in BOLD signal would only be weakly affected by task demands and could be separated when entangled with the task-related signals (Fair et al., 2007; Fox et al., 2006). Several studies have implemented this approach on task-based fMRI data to yield the intrinsic functional connectivity network (e.g. Bassett et al., 2011; Boets et al., 2013; Ebisch et al., 2013; Fair et al., 2007).

For each subject, a GLM was performed to assess bivariate Pearson correlation coefficients between ROIs’ BOLD time-series. These coefficients were averaged across subjects. As a result, networks of functionally connected (communicating) regions were uncovered. We refer to this clustering as the connectivity network.

### 2.6 Signal-to-noise ratio measurement

To measure the reliability of the fMRI signal for the activation (from UNIVAR), representation (from RSA) and connectivity (from FCA) network, we randomly split the participants into two groups (*n* = 10 or 11 per group). For each of these analyses, we correlated the resulting activation, representation, and connectivity network matrices (off-diagonal values) of one group with the matrix of the other group. This process was performed for a total of 100 iterations (each time randomly splitting the data into two groups). The correlations were adjusted with the Spearman-Brown split-half reliability formula and then averaged (across the 100 iterations) for UNIVAR, RSA, and FCA separately. The results from the between-subject correlations work as a measure of signal-to-noise ratio (SNR), taking the between-subject variability in the neural data into account, in that it estimates the maximum correlation we could expect. The correlation between the same types of data from the two sub-groups (group 1 vs. group 2 in FCA results) should be higher than the correlation with another type of data (e.g., FCA vs. RSA results). This SNR correlation coefficient was also squared to obtain the proportion of the variance in the signal that can be explained by other variables.

### 2.7 Anatomical proximity

For each ROI per hemisphere, we collected the x-y-z coordinates of its voxels. Consequently, for each ROI pair, we calculated Euclidian distances for all possible pairs of voxels between these two ROIs. Among these calculated distances, we use the minimum value per ROI pair as a measure of the anatomical distance between the two ROIs. Then, we averaged the distances across the two hemispheres. We also performed supplementary analyses with distance based on the average rather than the minimum value, which yielded very similar results (the two indices correlate strongly, *r* = .81). As a final step, we inverted these results to have a measure of anatomical proximity instead of distance with the maximal distance becoming the minimal proximity zero. We refer to these results as the anatomical proximity network. Dependency of functional connectivity on anatomical distance has been observed (Salvador et al., 2005). Thus, the anatomical proximity network was included in the partial correlation and the multiple regression model to rule out the effects of anatomical proximity when comparing the activation, representation and connectivity network.

### 2.8 Comparing the activation, representation and connectivity network

#### 2.8.1 (Partial) correlation models

To understand how similar the activation, representation, connectivity and anatomical proximity network are, we conducted a rank-order correlational analysis between these networks. In addition, we also computed the partial Spearman correlation coefficient to understand the similarities between the two networks while controlling for the remaining networks. To draw statistical inferences, we conducted the permutation test, wherein one of the variables of interest (one of the networks, consisting of all possible unique ROI pairs (120 pairs)) was randomly shuffled and then (partially) correlated with the unshuffled variables (remaining original networks, each consisting of all possible unique ROI pairs (120 pairs per network)). This process was iterated 1000 times. These permutation tests provide empirical *p*-values reflecting the proportion of permutations wherein the (partial) correlations with the shuffled data were larger (or equally large) than the original (partial) correlations.

#### 2.8.2 Multiple regression models

Following up on the (partial) correlation models, we conducted multiple regression analysis to investigate if the activation, representation or connectivity network respectively, could be explained by the other remaining networks. The anatomical proximity network was also included in all of the multiple regression models. Z-score standardizations were performed to normalize the data before building a regression equation. Similarly to the correlational analysis, permutation tests were used to obtain empirical *p*-values. In the end, the percentage variance explained by the model was compared to the squared signal-to-noise ratio of the predicted variables of the model.

#### 2.8.3 Multi-dimensional scaling (MDS) and Procrustes transformations

We conducted multidimensional scaling (MDS) on the activation, representation, and connectivity network matrices to visualize the networks in a two-dimensional space that shows the distance between each pair of ROIs based on how dissimilar these ROIs are in terms of their activation, representation, and connectivity respectively. MDS results of the representation network were used as a template to which the MDS results of the activation and connectivity networks were aligned using Procrustes transformations, to visualize the networks on the same space.

## 3. Results

### 3.1 Networks

In total, we have four matrices (see Fig. 3). For three of the methods (RSA, UNIVAR and FCA) the values in the matrices are based upon correlational analyses. In each of these matrices, we had a large range of values. For the representation network, for which vectorized first-order RSA results were correlated between all ROI pairs, correlations range from .07 (V5 – insula) to .82 (BA3 – BA4). In the activation network matrix, the correlation results range from -.01 (precuneus – PO) to .98 (BA3 – BA4). The values of the ROI-to-ROI connectivity range from -.17 (precuneus – PO) to .83 (BA3 – BA4). The anatomical proximity network values range from 0 to 67.63. The higher the value, the more closely the two ROIs are located. As the values are inverted distances, a value of 0 indicates the minimum anatomical proximity between ROIs (e.g., BA1 – BA17), which in the original distance was 67.63 mm. A proximity value of 67.63 indicates the maximum anatomical proximity between ROIs: these ROIs are located right next to each other (e.g., BA1 – BA2).

**Fig. 3.**
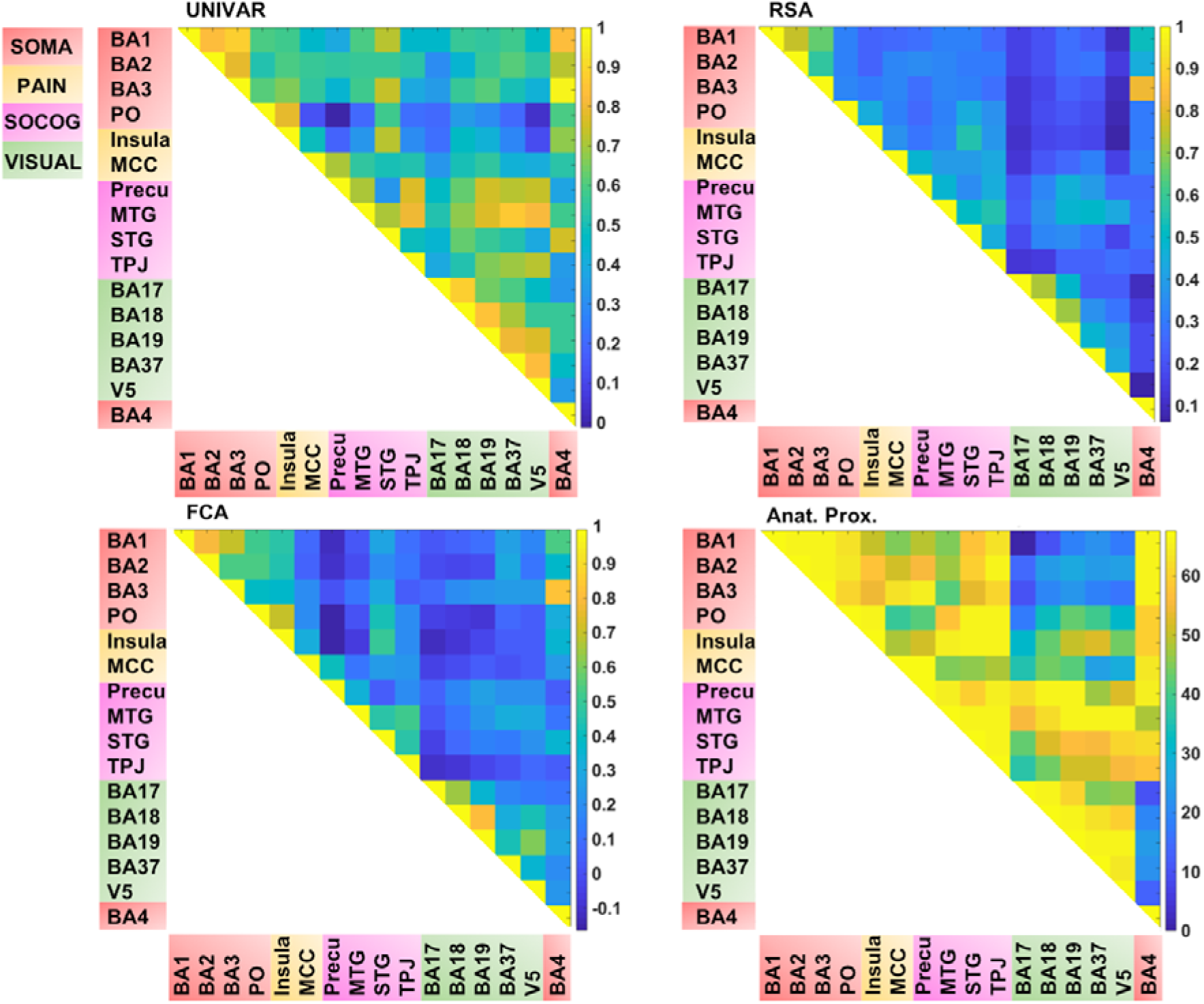
Visualization of the different networks before z-score standardization of the correlation coefficients. Top left: activation network (UNIVAR), top right: representation network (RSA), bottom left: connectivity network (FCA), bottom right: anatomical proximity network. Yellow in the matrix = when two ROIs are very similar in their activation (UNIVAR results) or their representation (RSA results), are well connected (FCA results), or are located closely in the brain. Blue in the matrix = when two ROIs are very different in their activation (UNIVAR results) or their representation (RSA results), are not connected (FCA results), or are located remotely in the brain. SOMA (red) = somatosensory-motor network areas, PAIN (yellow) = pain network areas, SOCOG (purple) = social-cognitive network areas, VISUAL (green) = visual network areas

Each of the matrices was very reliable. The signal-to-noise ratio estimated from the results of between-subjects correlations was *r* = 0.92 (squared to obtain explainable variance: EV = 85%) for the representation network, *r* = 0.96 (EV = 92%) for the activation network, and *r* = 0.97 (EV = 94%) for the connectivity network. As illustrated in Fig. 3, the representation, the activation, and connectivity networks look highly similar to each other. For example, the high correlation values between the ROIs in somatosensory areas such as BA3, BA1, and BA2 are apparent in all of these networks: these ROIs contain similar task-related information (based on RSA results), are activated to a similar level (based on UNIVAR results) and are functionally linked to each other (based on FCA results). BA4, the motor area, is strongly correlated to BA3 and BA1 in the representation, activation, and connectivity network, but only moderately to BA2. Another example is the moderate to high correlation between visual areas, found in the representation, activation and connectivity network. In sum, this finding applies to all the four ROI networks. Areas of different ROI networks typically show lower correlations, which is again consistent across methods. For example, the moderate correlations between social-cognitive areas and visual areas can be found in the representation, activation and connectivity network.

### 3.2 Comparing networks

To understand the (dis)similarity between representation, activation, connectivity, and anatomical proximity networks more quantitatively, we tested the linear relationship among these networks. The results indicated that all networks are similarly organized in the context of brain function and anatomy, with the Spearman rank-order correlations (all significant) ranging from .53 to .79 (see Table 1). In addition, the partial correlation coefficients were computed between two networks after removing the effect of the other remaining networks. The results from partial correlation (including all four networks) demonstrated that, after controlling for the other networks, the activation and connectivity network (*r*_*S*_ = .50, *p* < .001), the representation and activation network (*r*_*S*_ = .34, *p* < .001), and the representation and anatomical proximity network (*r*_*S*_ = .67, *p* < .001) still correlate significantly (see Table 2). Conversely, the measured partial correlation between the representation and connectivity network was no longer significant after ruling out the effects of the other covariates (partial *r*_*S*_= .10), implying that their association is fully explained by their relationship with other networks. The partial correlation between the connectivity and anatomical proximity network (partial *r*_*S*_ = .15), and between the activation and anatomical proximity network (partial *r*_*S*_ = .06) was also no longer significant.

**Table 1.** The correlations between the activation (UNIVAR), representation (RSA), connectivity (FCA) and anatomical proximity (Anat. Prox.) network. * = significant correlation.

|  | UNIVAR | RSA | FCA | Anat. Prox. |
| --- | --- | --- | --- | --- |
| UNIVAR | 1 | .66* | .70* | .53* |
| RSA | .66* | 1 | .61* | .79* |
| FCA | .70* | .61* | 1 | .55* |
| Anat. Prox. | .53* | .79* | .55* | 1 |

**Table 2.** The partial correlations between the activation (UNIVAR), representation (RSA), connectivity (FCA) and anatomical proximity (Anat. Prox.) network. * = significant partial correlation.

|  | UNIVAR | RSA | FCA | Anat. Prox. |
| --- | --- | --- | --- | --- |
| UNIVAR | 1 | .34* | .50* | -.10 |
| **RSA** | .34* | 1 | .10 | .67* |
| **FCA** | .50* | .10 | 1 | .15 |
| **Anat.**<br>**Prox.** | -.10 | .67* | .15 | 1 |

As an alternative approach, we also implemented multiple regression models. Similar to the (partial) correlation measurements, these regression models quantify the relations between the networks, but in addition the regression models provide an estimate of the total variance in a network that can be explained by all other networks.

A first model tested if the connectivity, activation and anatomical proximity network significantly predicted the *representation* network. The coefficient of determination from the regression equation indicated that these three predictors explained 71.2% of variability in the representation network (*R*^*2*^ = .712, *F*(3,116) = 96, *p* < .001). The squared signal-to-noise ratio (based on the between-subjects correlation) in the representation network indicated 85% of the variance to be explainable, leaving approximately 14% of the signal unexplained. In addition, we calculated the *β* coefficients to examine the degree to which each predictor independently contributes to the prediction of the representation network. According to the results, the anatomical proximity network significantly contributed to the prediction of the representation network (*β* = 0.40, *p* < .001), as did the connectivity network (*β* = 0.36 *p* = .004) and the activation network (*β* = 0.26, *p* = .03).

Similarly, we predicted the *connectivity* network based on the representation, activation, and anatomical proximity network, using multiple regression analysis. The results indicated that the predictors explained 59.6% of variability in the connectivity network (*R*^*2*^ = .596, *F*(3,116) = 57, *p* < .001). The squared signal-to-noise ratio (based on the between-subjects correlation) in the connectivity network indicated 94% of the variance to be explainable, leaving approximately 34% of the signal unexplained. When examining the independent contributions of each predictor, we found out that the representation network significantly contributed to the prediction of the connectivity network (*β* = 0.51, *p* = .003), as did the activation network (*β* = 0.37, *p* = .005), but not the anatomical proximity network (*β* = −0.07, *p* = .55).

Lastly, we tested if the representation, connectivity, and anatomical proximity network significantly predicted the *activation* network. The results revealed that the predictors explained 55.8% of variability in the activation network (*R*^*2*^ = .558, *F*(3,116) = 49, *p* < .001). The squared signal-to-noise ratio (based on the between-subjects correlation) in the activation network indicated 92% of the variance to be explainable, leaving approximately 36% of the signal unexplained. The predictors indicated that the representation network significantly contributed to the prediction of the activation network (*β* = 0.41, *p* = .01), as did the connectivity network (*β* = 0.41, *p* = .001), but not the anatomical proximity network (*β* = −0.01, *p* = .91).

Thus, for each type of network, we find that a lot of the structure can be predicted from the other networks, but there is also some remaining variance left unexplained. We visualized this unique signal left in each of these networks after regressing out the signal explained by the other networks from the representation, activation and connectivity network respectively (see Fig. 4b). In Fig. 4b, in contrast to Fig. 3 and 4a (which takes the values of Fig. 3 and z-score standardizes them, for reasons mentioned above), the networks now do not look similar: they show different patterns.

**Fig. 4.**
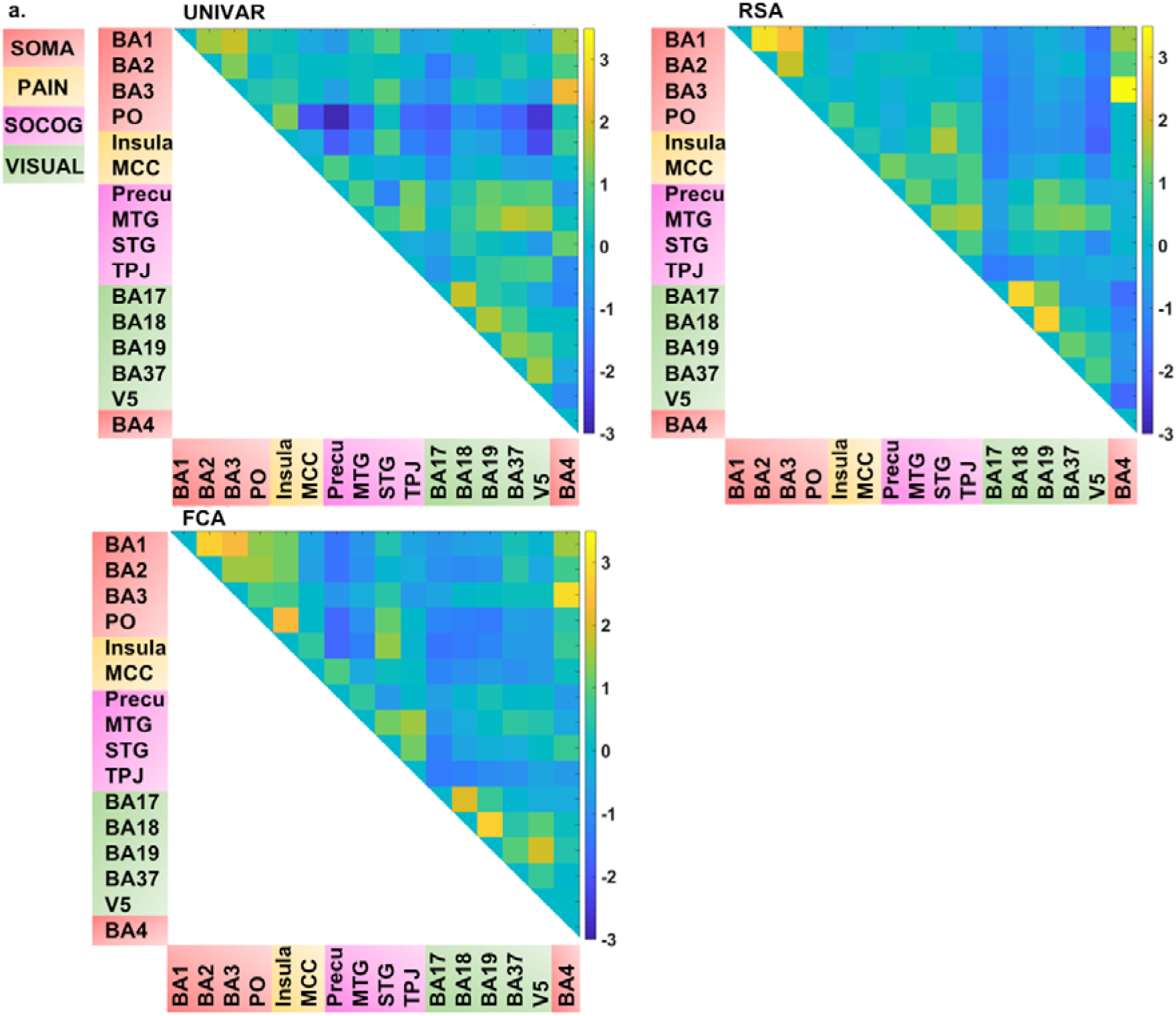
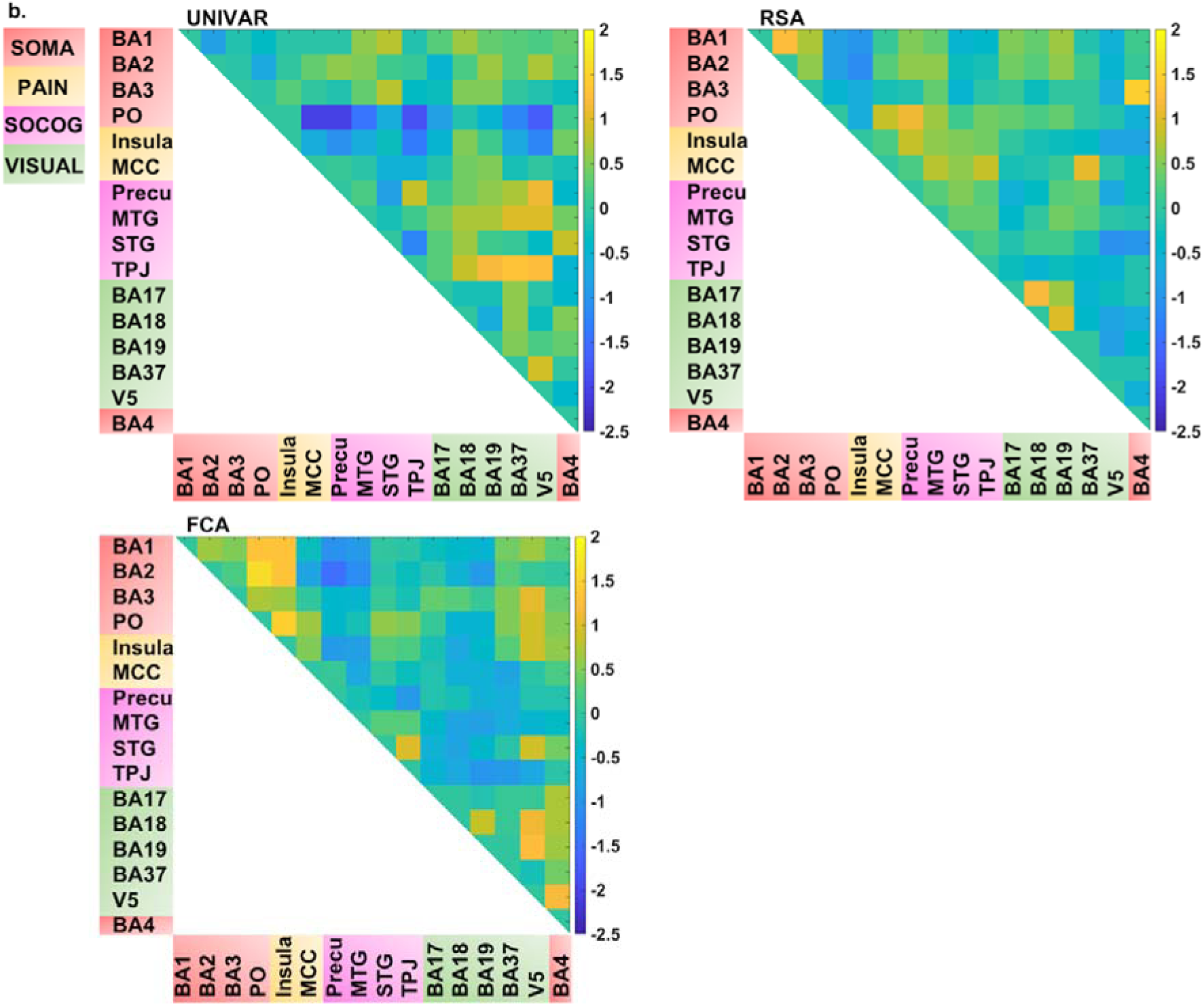
**a.** Visualization of the different networks after z-score standardization of the correlation coefficients. **b.** Visualization of the different networks after regressing out the signal explained by the other networks. **In a. and b.** Top left: activation network (UNIVAR), top right: representation network (RSA), bottom left: connectivity network (FCA), bottom right: anatomical proximity network. Yellow in the matrix = when two ROIs are very similar in their activation (UNIVAR results) or their representation (RSA results), are well connected (FCA results), or are located closely in the brain. Blue in the matrix = when two ROIs are very different in their activation (UNIVAR results) or their representation (RSA results), are not connected (FCA results), or are located remotely in the brain. SOMA (red) = somatosensory-motor network areas, PAIN (yellow) = pain network areas, SOCOG (purple) = social-cognitive network areas, VISUAL (green) = visual network areas

Several unique findings concerning correlations between ROI-networks can be observed in Fig. 4b. For example, social-cognitive brain areas correlate strongly to other visual areas in the activation network (e.g., *r* (before z-score standardization) = .69 between TPJ and BA37) while this is moderate to low in the representational (e.g., *r* = .24 between TPJ and BA37) and connectivity network (e.g., *r* = .01 between TPJ and BA37). This finding implies that these areas are activated similarly, but do not represent similar information nor do they communicate with each other. Another example, social-cognitive areas correlate moderately to somatosensory-motor areas (e.g., *r* = .32 (representation), *r* = .59 (activation) between MTG and BA1), except in the connectivity network (e.g., *r* = .03 between MTG and BA1). As a last example, visual area V5 shows a moderate correlation to other brain areas in the representation network (e.g., *r* = .39 between V5 and BA19) while a much stronger correlation is found in the other networks (e.g., *r* = .76 (activation) *r* = .61 (connectivity) between V5 and BA19).

For visualization purposes, we performed MDS on the three types of dissimilarity matrices to reconstruct two-dimensional spatial configuration that reflects the proximity in the matrices. Moreover, Procrustes transformations were performed to align the configurations. The resulting configurations are shown in Fig. 5. The results confirm the high similarity (*d* (Procrustes distance: the difference between the shape of the two networks) between the activation and representation network = .48, *d* between the connectivity and representation network = .34, *d* between the activation and connectivity network = .42) and some dissimilarities between the networks as was previously indicated by the (partial) correlation and multiple regression models. As an example of correspondence between the three networks, Fig. 5 shows that somatosensory-motor areas are located nearby in all three networks, implying high similarity in activation and representation and strong inter-regional communication among these areas. As an example of a difference between the networks, the social-cognitive brain areas are places close to visual areas overall in the activation network (blue in Fig. 5) but not so much in the other networks. It suggests that social cognitive brain areas and the visual cortex do not represent the same information and that those areas are not functionally connected despite the similar magnitude of neural response.

**Fig. 5.**
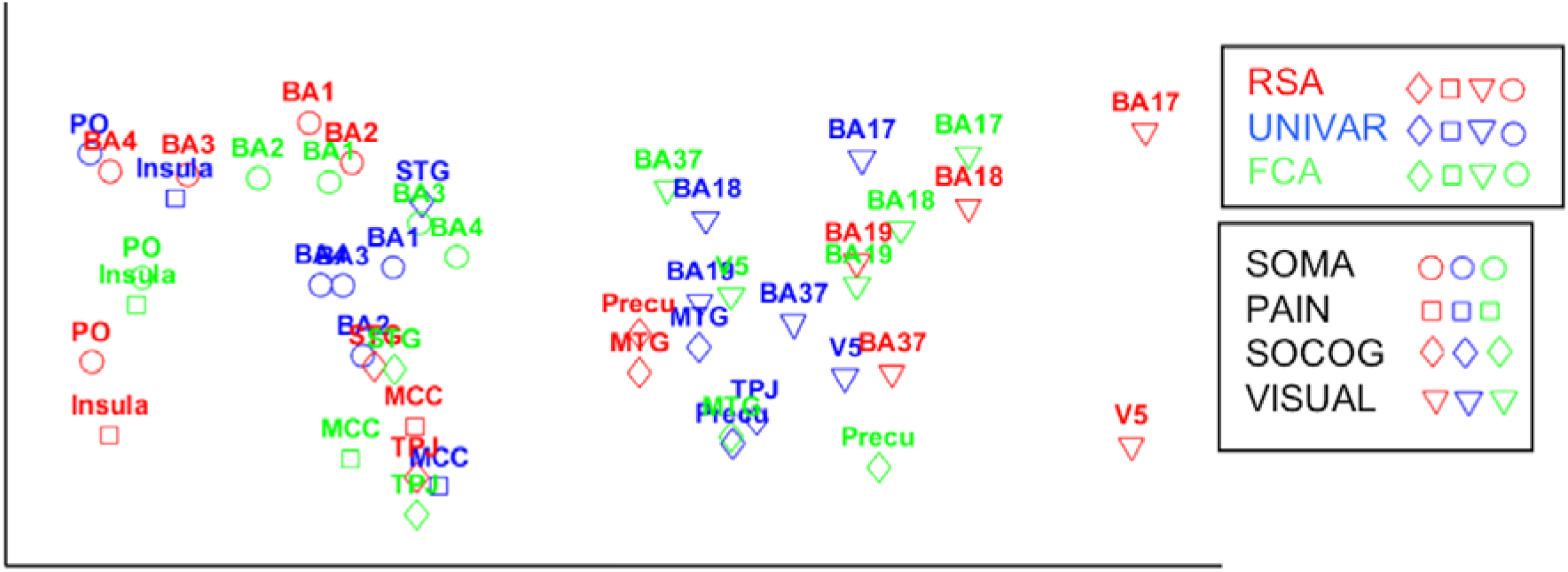
Procrustes transformed MDS results of the activation (UNIVAR, blue) and connectivity network (FCA, green) to the MDS results of the representation network (RSA, red). SOMA (circles) = somatosensory-motor network areas, PAIN (squares) = pain network areas, SOCOG (diamonds) = social-cognitive network areas, VISUAL (triangles) = visual network areas

## 4. Discussion

RSA has recently emerged as a method for investigating how brain regions are organized into networks. UNIVAR and FCA are two other popular methods for analyzing fMRI data to understand the functional structure of brain networks. Although two or more of these methods have been used simultaneously to analyze the same set of data in many studies, most of them have focused on the properties of each ROI separately. No study, to our knowledge, directly and simultaneously compared networks derived from RSA with networks built with UNIVAR and FCA. In the current study, we explored how the structure of networks built from RSA, UNIVAR and FCA relate to each other after ruling out the effect of the anatomical location of network nodes (ROIs). We analyzed fMRI data of a previous study (Lee Masson et al., 2018) with these methods and performed (partial) correlation and multiple regression analysis on the resulting networks.

The current study reveals that neural networks resulting from RSA, UNIVAR and FCA are highly similar even after ruling out the effect of anatomical proximity. As predicted, brain areas within the somatosensory-motor network are similarly activated, represent similar task-related information, and are intrinsically connected. This applies also to the other sub-networks (pain, social-cognitive and visual). As outlined in the introduction, RSA, UNIVAR and FCA share theoretical and/or methodological properties that can explain similarities as observed in this study. The high similarity in the neural networks of RSA and FCA provides support for the idea that brain areas showing similar stimulus-related selectivity are also intrinsically connected. Our finding is in line with previous resting-state fMRI studies that have identified functionally relevant networks, such as the primary visual network, auditory network, motor network, and cognitive networks, during rest (e.g., Biswal et al., 1995; Fox and Raichle, 2007; Jung et al., 2018).

On the other hand, our finding suggests that the network structure derived from each method contains unique signals. To reveal this, the explainable variance of each network revealed by SNR estimation was compared with the actual variance explained by the other networks. These results suggested that the network, derived from each method, contain idiosyncratic structure that none of the other networks are able to explain. Analyzing the remaining signal variance that was left unexplained, we were also able to reveal the idiosyncratic network structure of each method. For example, brain areas in the social-cognitive network are similar to areas in the visual network in terms of neural activation, whereas neural patterns of those two sub-networks do not represent the same information and they are not intrinsically connected. Another example are the moderate correlations between social-cognitive brain areas and somatosensory-motor areas in the activation and representation network, but not in the connectivity network.

This idiosyncratic structure is important to keep in mind when interpreting a network structure found with one particular method. Although second-order RSA can be used to construct brain connectivity, RSA and FCA adopt different approaches shown in their methodology: correlating RDMs in RSA; correlating the BOLD signal fluctuations in FCA. Thus, RSA is used for investigating the similarity between brain areas in how they represent the task-related information while FCA is used for investigating how a series of brain areas construct the intrinsically connected cortical network. These distinctions allow RSA and FCA to tap into the functional architecture of the brain from different perspectives as revealed in the idiosyncratic network structures.

Likewise, the same reasoning can be applied to the relationship between UNIVAR and FCA, and RSA and UNIVAR. As outlined in the introduction, they are related theoretically and empirically while they differ in their focus, allowing both similarities and dissimilarities between the resulting networks.

Such distinctions between the network structures derived from different methods has also been observed in the recent study of Jung and her colleagues (Jung et al., 2018) comparing resting-state fMRI and structural connectivity. Although their comparison involves different methods than ours, they provided some possible explanations that should be considered in the current study. Quality and nature of the datasets used for three methods (even from identical data sources, but measured at different times or analyzed in a different way) may not be equal and different measurement noise may be present (Jung et al., 2018). In addition, they mention that networks during mental activity are modulated away (slightly) from intrinsic connections, which is especially relevant to the comparison of RSA with FCA. Accordingly, our findings of similarities and differences between RSA and FCA network structure are consistent with the observation that studies using both RSA and FCA lead to either similar or different conclusions about brain function derived from the two methods (e.g., Boets et al., 2013; Zeharia et al., 2015). Despite the high similarity across the network structures derived from the UNIVAR, RSA and FCA methods, given the nature of idiosyncrasy of each network, we encourage researchers to understand the benefits of each methodology and what they (do not) detect; and to use them adequately depending on the research questions.

As a critical note, we point to several limitations of our current study. First, the current findings are based on only one task domain (i.e., social touch videos perception), and our conclusions should be complemented by future studies that include other tasks, such as moral decision-making tasks, or tasks using other sensory modalities such as auditory and tactile scenes. RSA and UNIVAR methods may not produce similar network structures in another task. This could in particular be the case, when having a task with no activation differences across the conditions but evoking neural pattern selectivity. Second, we selected a limited number of ROIs rather than including a large number of network nodes. One important argument for doing this is that the selected brain regions had to include meaningful task-related signals for performing RSA (see the description of diagonal versus non-diagonal measures as a reliability test in choosing ROIs in Methods). The effect of the number and size of ROIs on the relationships between the networks obtained using RSA, UNIVAR and FCA can be explored further. Finally, extending the comparisons made in the current study is another important step to take. Specifically, networks built from second-order RSA and multivariate functional connectivity could also be compared (Anzellotti and Coutanche, 2018; Coutanche and Thompson-Schill, 2014).

## 5. Conclusions

The present study provides first-time evidence that cortical network structures derived from three commonly used neuroimaging approaches (RSA, UNIVAR and FCA) are highly similar regardless of the structural variations of each network. Importantly, the study also demonstrates that each of these three networks contains idiosyncratic structure, unexplainable by the other networks. As such, all three methods are important when investigating the functional organization of networks in the brain. Improving the understanding of the relationship between the structures of the networks derived from these methods will allow researchers to use RSA, UNIVAR and FCA more adequately.

## Data availability statement

The data for this study can be found in the Open Science Framework (https://osf.io/b4np9/).

## Abbreviations

MVPA: multi-voxel pattern analysis;
RSA: representational similarity analysis;
UNIVAR: univariate analysis;
FCA: functional connectivity analysis

## Author contributions

I.P., H.O.d.B., and H.L.M. set up the project. FMRI data collection: see Lee Masson et al., 2018. I.P. and H.L.M. analyzed the data. I.P., H.O.d.B., and H.L.M. wrote the paper.

